# Convection-enhanced diffusion and directed withdrawal of methylene blue in agarose hydrogel using finite element analyses

**DOI:** 10.1101/2022.10.27.514109

**Authors:** C. Shaw, K. Hossain, Cecile Riviere-Cazaux, Terry Burns, M. Rashed Khan

## Abstract

Precision drug delivery for optimized therapeutic targeting requires knowledge of momentum transport and molecular diffusion of molecules within the patient’s interstitial tissue, especially for tumor treatment within the brain. Dispersion in the interstitial space is impacted by delivery method, tissue material properties, individual-specific fluid flow, and particle size of the input solute. Knowledge of a drug’s dispersion allows for optimizing solute delivery, concentration, and flow rates to maximize drug distribution and biomarker recovery. For delivering drugs, increased knowledge of drug location after delivery can improve therapeutic treatment by optimizing the dosing of healthy and unhealthy tissue. Finite element methods (FEM) tools, such as COMSOL Multiphysics, can simulate molecular distribution inside-individual specific shapes and porous material properties. Furthermore, an additional unmet need is delivery methods that can be adjusted to manipulate diffusion regions through tissue via techniques such as directed flow. This would be especially valuable in targeted drug delivery within tumors to increase the cancerous surface area covered while limiting damage to surrounding tissues. In this project, the directed flow was induced by perfusing the injected solution at an input probe while withdrawing fluid at an output probe, enabling targeted flow through the desired region. FEM computation faithfully replicated these conditions and could be used to determine the effective concentrations perfused over the region of interest. We leveraged COMSOL Multiphysics to perform a computational study simulating convection-enhanced delivery (CED) with an output probe pulling the concentration profile over the region of interest. This simulation system can be applied to therapeutics targeting, vaccine subcutaneous injection, and waste and media diffusion in tissue engineering.

## 1. Introduction

### 1.1 Clinical Background

Primary and metastatic brain tumors are central nervous system (CNS) diseases characterized by high disability and mortality and limited curative treatment options [1]. Several modalities have emerged as cancer treatment options, including chemotherapy[2], immunotherapy[3], and cell-type specific targeted therapy[4]. Despite these advancements, delivering these therapeutics past the blood-brain barrier (BBB) and into the CNS remains a significant challenge. Osmotic opening and ultrasound-induced BBB disruption have emerged as two new opportunities to increase drug delivery. However, the effects of these methods are transient and heterogenous across and within patients. Alternatively, convection-enhanced delivery (CED) has emerged as a technique to increase local drug delivery into brain tumors[5]. In CED, the interstitial infusion of an agent by a syringe pump is able to create a pressure gradient, permitting enhanced distribution of CNS therapeutics throughout the tumor following focal delivery of an agent.

CED with an output probe extracting solution can perturb solute diffusion in porous media[6]. In CED, a solute is injected through a probe into porous media to induce fluid convection and to allow greater solute diffusion through the media via a pressure differential. Porous media describes a solid material containing gaps filled with fluid. The dispersal of a solute inside a porous media is impacted by diffusion of the solute in the liquid and interaction of the solute and liquid with the matrix. With fluid convection being an additional component to diffusing solutes, increased control to diffusion allows improved control of solute dispersal. Solution convection and diffusion in a porous media describes systems such as drug delivery into tissue[7] and biomolecule motion within tissue[8].owever, optimization of CED to improve drug delivery has been limited by an inability to determine the extent of drug diffusion and relative therapeutic impact based on biomarker recovery. Consequently, modeling of CED and evaluating the resulting indications of solute spread can improve patient outcomes.

### 1.2 Software and Physic Background

COMSOL Multiphysics (Version 5.6, Burlington, MA) is a software package that conducts finite element method solutions of differential equations to converge to a solution of the equations for given input boundary conditions. COMSOL Multiphysics can describe solution spread in a porous media. Modeling diffusion with these differential and algebraic equations includes descriptions of how diffusion and fluid flow occur and how fluid flow interacts with a porous matrix. As a solute moves within a medium, it exhibits Brownian motion, with its next direction of motion being a random walk. The particle in its initial condition will have equally sized vectors pointing in all directions around it. This is best described as Stoke-Einstein behavior[9]. The concentration gradient vector also impacts particle motion. In the presence of particles in a medium, a particle’s equally distributed random walk vectors will cause it to collide with other particles, particularly with increased collisions on the side with high concentrations-for our case, the side facing the input probe. Upon colliding with these particles, the net vector direction will be in the direction away from the input probe. Fick’s Second Law of diffusion best describes this net vector.

### 1.3 Previous Work

Different tiers of CED modeling have been demonstrated. One model utilized ANSYS FLUENT (ANSYS Inc., Canonsburg, USA) to simulate injection into a liquid modeled as a 2 dimensional plane of an eye[10]. Rather than model fluid momentum from a probe, this article placed solute inside the eye and added laser heating to induce fluid convection in the eye. Another tier of modeling (using COMSOL Multiphysics) focused on diffusion with variables other than fluid convection. Some setups include diffusion in porous media with temperature-induced variation in diffusivity[11], diffusion with a chemical reaction[12], diffusion in a porous media in an electric field[13], diffusion of solute amongst two fluids with one displacing the other[14], and diffusion in a liquid in a tube replicating nerve fibers containing semipermeable membranes for walls[15]. Furthermore, simulations of brain characteristics have been demonstrated with a probe compressing brain tissue[16], changes in chemical reactions with motion induced by probes[17], and a brain model with tumor growth represented as an advancing concentration profile of tumoral cellular spread[18]. Two models using COMSOL Multiphysics simulated injection into skin and tumors. The skin injection model evaluated solution motion after injection, albeit without momentum ignoring more rigorous considerations[19]. This model evaluated tissue deformation following solute injection and its impact thereof on porosity and fluid flow, each affecting solute diffusion, advection, and adsorption. Another model focused on drug transport within a generalized tumor, evaluating tumor conditions such as tissue permeability that vary-interstitial flow and pressure, as identified by Baxter and Jain[20][21]. While this simulation did not evaluate the impact of solute injections on particle flow and diffusion, it did note changes in drug motion for both varying sizes and shapes of tumors and differences in pressure and fluid flow. More rigorous models have simulated solute diffusion and additional fluid convection that is following present injection into a brain. One model using Ansys Fluent injected the solution into a simple brain model to evaluate the differences amongst diffusion only, convection-enhanced delivery from a single probe, and delivery from a multi-holed probe[22]. Other efforts have been made to simulate CED, including treating glioblastomas (GBM)[23]–[26]. Rosenbluth et al. and Sampson et al. utilized a software package based on partial differential equations and boundary conditions developed by Morrison et al. and marketed ‘Therataxis, Baltimore, MD, USA; and BrainLAB, Feldkirchen’[23], [24], [27]. Using Ansys Fluent, Zhan et al. has developed multiple CED models of injection into brains[25], [26], [28]. These models primarily focused on adjusting interstitial fluid flow (IFF), diffusivity corresponding to specific drugs, and probe placement near a tumor. However, review by Stine et al. indicates that Rosenbluth et al. and Sampson et al. mathematical models have failed to improve patients’ outcomes for the application in drug delivery. Furthermore, these CED simulations only modeled a single injecting probe diffusing a sphere of solute around the outlet without creating the conveyor belt of solute induced with an output probe.

### 1.4 Improved CED Simulation

CED conventionally only includes an input probe for injection; however, in this simulation, an output probe is utilized to induce a “conveyor belt” of flow from the input probe to the output probe, driven by an increased volume of inflow liquid, pressure, and solute neighboring an outflow region acting as a sink via local solute depletion. This computational model allows for the analysis and optimization of the technique of CED. We can precisely determine the resulting concentration profile in space, indicating the dose within tumorous and healthy tissue. Hydrogels are porous media composed of solid (polymer) and liquid (water) materials. Agarose hydrogels have been applied for use as a model of brain and tumor tissue [7] [33] due to their similarities in diffusion, permeability, and mechanical properties. With agarose replicating brain tissue properties, using the *in-silico* model to duplicate agarose *in vitro*, creates an efficient validation technique.

In addition to diffusion and fluid flow, and the interaction of the latter with a porous matrix, our work has added fluid convection as another vector influencing particle motion. We do this by modeling fluid injection into a hydrogel through a probe followed by fluid extraction in a secondary nearby probe. This induces fluid momentum transfer through the gel and mass diffusion of molecules into the gel matrix. In the Navier-Stokes equation, the 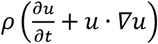 term relates convection with fluid velocity and density to yield fluid momentum. However, momentum in this equation only corresponds to the transfer of momentum from mass to mass. To describe how a particle moves in the fluid, we utilize the convection diffusion equation to include the contribution of the convection vector to the net vector influencing particle motion. By evaluating these three vectors—random walk, concentration gradient, and fluid convection -- we can determine the net force on particles in a medium and how this affects particle dispersion.

As can be noted, there are multiple gaps in successfully utilizing convection enhanced delivery and in COMSOL Multiphysics simulations covering fluid flow and diffusion. Therefore, we have conducted simulations to bridge these gaps. We designed an *in vitro* test replicating geometry to induce convection enhanced delivery in the presence of an output probe to induce controlled momentum transfer and mass diffusion within COMSOL Multiphysics. We aim to establish an enhanced CED computational framework to demonstrate, measure, and optimize perturbed molecular diffusion for use in drug delivery planning for optimized dosing of tumors

## 2. Materials and Methods

### 2.1 Modelling and Simulation

#### 2.1.1 Governing Equations

COMSOL Multiphysics takes differential equations and arrives at an approximate numerical solution through finite element methods [38]. Three sets of differential equations are utilized, including the Navier-Stokes Equations for laminar flow through the probes[39], the Brinkman Porous Flow Equations for flow through the porous hydrogel[14], and Fick’s Second Law of Diffusion for diffusion of the solute listed as Equation 1, Equation 2, and Equation 3 respectively[39], [40]. Equation 4, shows the Convection Diffusion equation COMSOL Multiphysics uses to combine fluid flow and diffusion[41]. Within the equations, the terms cover fluid velocity (μ), fluid pressure (p), fluid density (ρ), fluid dynamic viscosity (μ), external applied forces (F), time (t), stress tensor (T), identity vector (I), porosity (ε), diffusion flux (J), coefficient of diffusion (D), forchheimer drag coefficient (β_F_), and Brinkman Flow (Q_Br_), and concentration (c)[9], [14], [39], [41].

<u>Navier-Stokes Equation</u>

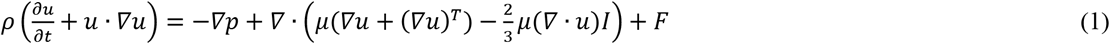

<u>Brinkman Porous Flow Equation</u>

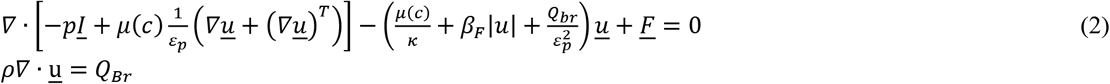

<u>Fick’s Second Law of Diffusion</u>

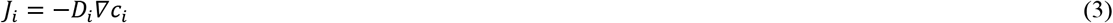

<u>Convection Diffusion Equation</u>

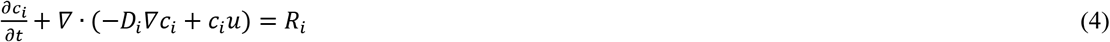

There is considerable difficulty in arriving at exact solutions for each equation. To circumvent this difficulty, FEM analysis arrives at approximate solutions that require boundary conditions to determine a constant value for an element.

#### 2.1.2 Model Geometry and Boundary Conditions

The boundary conditions of the differential equations are set in COMSOL by establishing the geometry of the system. In Figure 2A, we established the geometry to replicate a petri dish containing hydrogel with the probes embedded through the bottom of the petri dish. The open port tips of the probes opened midway through the gel. The walls of the probes replicated the in vitro setup by establishing the probe walls to have a no-slip boundary condition. The probes’ sizes, placement, and length, the petri dish dimensions; and the hydrogels properties are described in Table 1. Since it was not a part of the fluid flow or diffusion, the petri dish formed the barrier of the system but was not included in the computation. Meanwhile, the petri dish walls in contact with the fluid and porous matrix were defined to have no flux and a no-slip boundary condition (Fig. 2A). The top surface of the hydrogel was set as an open boundary to allow flux to occur. Initial conditions include injected solute having zero concentration in the gel and output probe. The input probe and upstream solute concentration was selected based on visual contrast between a known methylene blue input concentration followed by perfusion and diffusion in agarose hydrogel. Variables such as temperature, probe motion, and liquid type were kept constant throughout the simulations. The hydrogel geometry was set to replicate the interior volume of 60 X 15 mm petri dishes and the material properties were set to replicate 2% agarose hydrogel. The probe interior and the upstream fluid were set as water. Figure 2B shows the computational meshing that COMSOL constructed for fluid flow conditions. Analysis of the diffusion profile was conducted over the time domain represented between Figure 2C and 2E seen the probe tip. Data was extracted on linescans at the probe tip shown as the red line showing the probe length (Fig. 2G). The data from the line-scans resulted in the corresponding plots allowing for improved comparison (Fig. 2D, 2F, and 2H).

**Figure 1.**
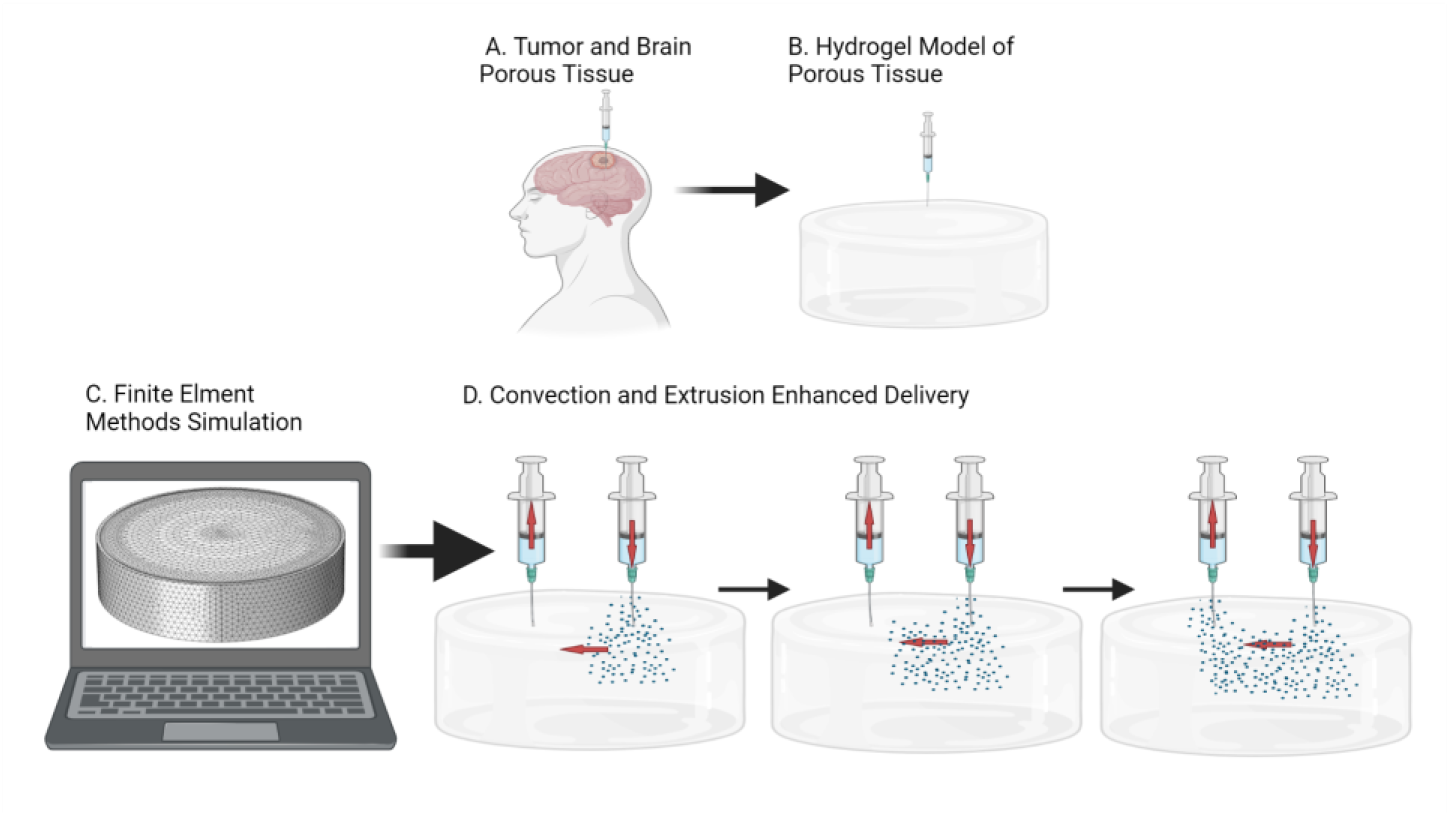
Simulation of diffusion. Convection enhanced delivery is a process of delivering solutes into porous matrices. (A) The brain’s diffusion properties can be mimicked by hydrogels (B) allowing *in situ* and *in silico* pairing tests. (C) Finite element methods can simulate convection and diffusion in porous matrices. (C) Injection of a solute and liquid induces diffusion and fluid convection in a porous media in conjunction with an extrusion probe. The two probes (one input and one output) induce a conveyor belt of fluid flow carrying the solute between them. Here we model convection enhanced delivery using COMSOL Multiphysics a finite element method software package.

**Figure 2.**
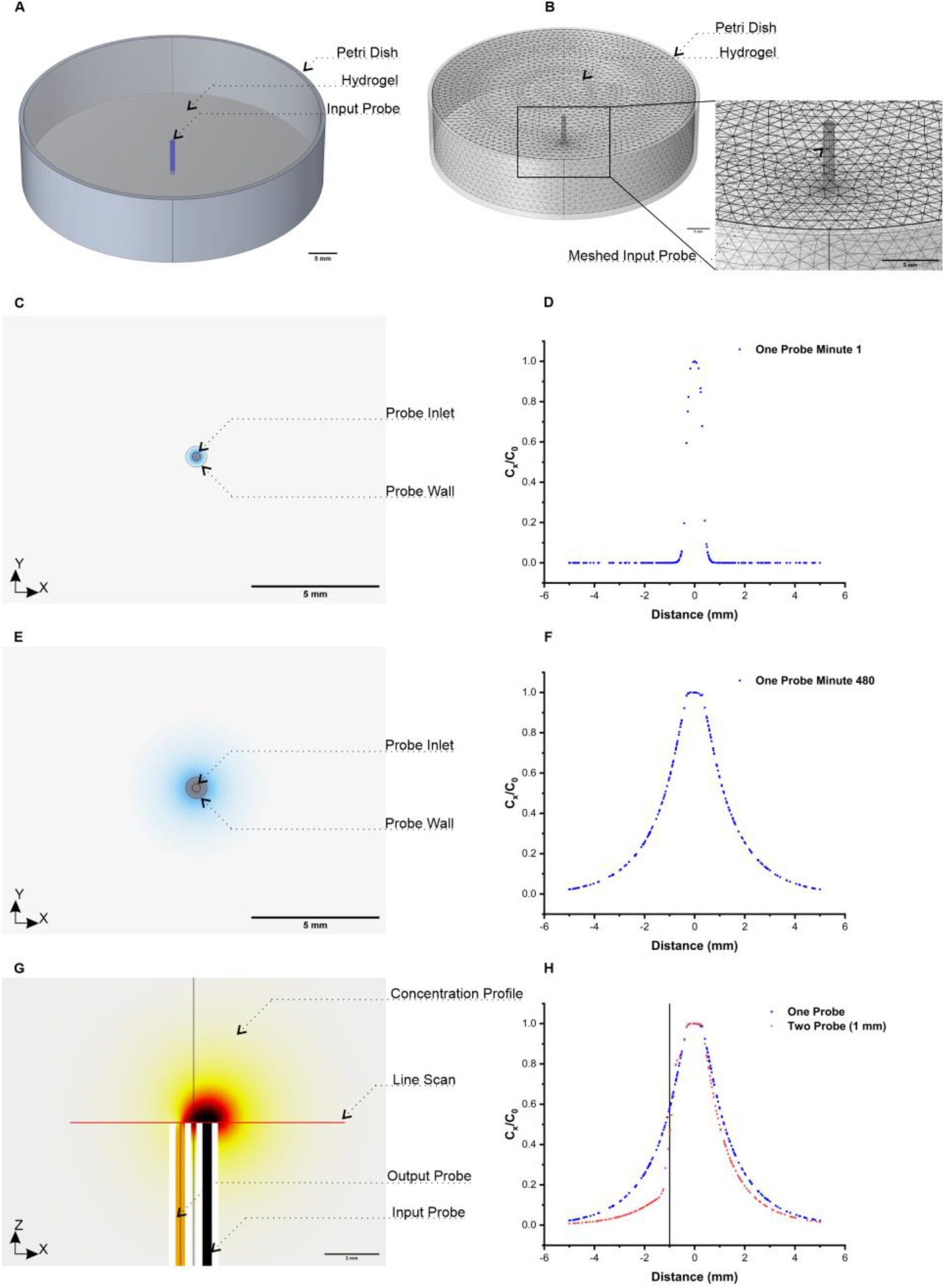
Finite element methods setup and continuous infusion. (A) illustrates the geometric boundary conditions implemented corresponding to a hydrogel sample in a petri dish with the probe in the center entering the gel from the base. (B) illustrates the meshing used by the finite element methods. The meshing is a portion of finite element methods to approximate a solution of differential equations solving for fluid flow and diffusion in the hydrogel. (C and E) are spatial illustrations of diffusion over time from the probe while (E and F) show the normalized concentration profiles centered on the probe outlet derived from line plots. (G) demonstrates a modified version of CED including an extrusion probe showing a perturb diffusion profile. (H) illustrates the normalized perturbation in comparison to the single probe diffusion profile.

**Table 1.**
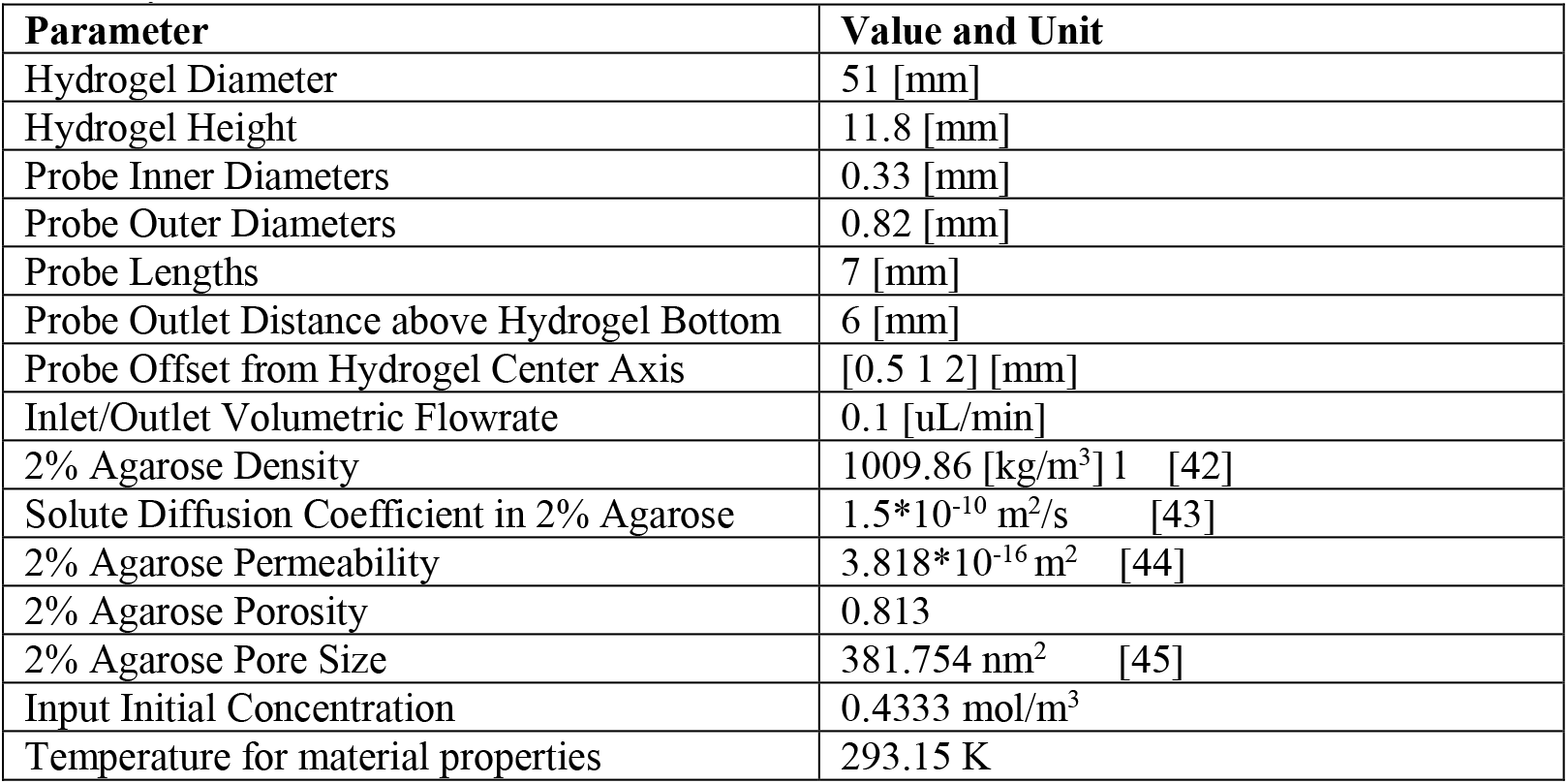
Perfusion Simulation Parameters.

#### 2.1.4 Mesh Generation and Simulation

The finite element method component of COMSOL creates a mesh to be solved. The mesh comprises of discrete spaces for which dependent variables in the differential equations can be estimated. The resolution of these meshes impacts the rigor of the simulation. Finer meshing consumes more time and requires greater computing power to complete. Larger meshing can be used to increase computational efficiency if the solution is converged. Since the meshed results are approximations over space, large spaces may not resolve finer details. Convergence is observed when the computed results do not vary with decreasing meshing sizes. To optimize our meshing, we determined the largest meshing size within convergence by conducting a series of simulations varying meshing size and elements. As seen in Supplemental Information, Figure S 1, “Fine” and “Finer” meshing results in convergence. Subsequently, using the “Fine” mesh to conduct the simulations, we compiled COMSOL to construct the meshes shown in Figure 2B with further meshing details listed in Table S1. Briefly, the amount of tetrahedrons in the simulation varied from 350,000 to 750,000 with the maximum and minimum size elements remaining consistent throughout the two probe simulations.

### 2.2 Experimental Validation

#### 2.2.1 Experimental Setup

A Model NanoJet pump was utilized to regulate pump headers (Chemyx Inc., Stafford, TX; Serial # 48011) and to compress and expand Polytetrafluoroethylene (PTFE) Luer Lock 1.0 mL glass syringes (Hamilton Syringe, Reno, NV; Part/Ref # 81320) which contained the methylene blue solution in order to inject into hydrogel. The output syringe contained dH_z_O to remove bubbles that prevent fluid extraction from the hydrogel. The syringes tipped with Luer lock blunt tip needles were connected to the gel via the PTFE tubing. The end of the PTFE tubing was placed within the 2% agarose hydrogel matrix (VWR Life Science, Radnor, Pa; CAS: 9012-36-6). The hydrogel was formed inside of 60 x 15 mm polystyrene petri dishes (VWR, Radnor, Pa; CAT: 25384-342) on the stereoscope containing a Mu1000-HS microscope digital camera (AmScope, Irvine, CA). The camera took an image once a minute for 8 hours while the injection and retraction flowrates were set to 0.1 uL/min. The injection solution had a concentration of 0.4333 mol/m^3^ of methylene blue (BioPharm, Hatfield, AR; 1% w/v, SKU: BM4292B) while the retraction syringe and tubing were primed with dH_2_O. Furthermore, we prepared the dye solution to 0.4333 mol/m^3^ by adding 420 μL of the 1% w/v methylene blue into 29.88 mL of dH2O. Images were taken by the AmScope and processed with ImageJ Fiji by National Institutes of Health (NIH) (Bethesda, MD).

For post-experimental processing, the data produced from ImageJ line plots were normalized across the probes as shown in the Supplemental Information Figure S 2 for comparison to simulation results. We imaged a digital caliper (Neiko Tools USA; NEIKO 01407A) to provide the ImageJ Fiji scale bar. The ImageJ Fiji Line Plot option provided pixel intensity on an 8-bit grayscale. The line plot was drawn to be 10 mm long, with the center on the input probe center and a second point over the output probe. Similarly, a line scan option in COMSOL Multiphysics yielded the simulated concentration over a similarly placed line located at the probe tip within the hydrogel. To normalize these data, the minimum value of the data set was subtracted from each value in the entire data set because the grayscale intensity values placed the relatively dark dyed regions close to zero while the light regions were close to 255. After this step, the data were then divided by the magnitude of the minimum value from the new data set. Adding this last data set by nominal gave us a normalized profile following the expected error function decay. Therefore, the COMSOL normalized concentration profile and the ImageJ grayscale value can be compared.

#### 2.2.2 Preparation of Agarose Hydrogels

The agarose polymers were weighed on a balance (VWR, Radnor, Pa; Model: VWR-314AC) and mixed with a solution of dH2O to a 2% (w/v) solution. The agarose and dH2O solution were placed on a heated stir plate (VWR, Radnor, Pa; CAT: No97042-714) at 125 °C for 15 minutes while covered and stirred continuously at 100 rpm to ensure complete dissolution. The gel was then placed in a 60 x 15 mm petri dish at 21 °C until it was completely solidified.

## 3. Results

### 3.1 Hydrogel Property Estimates

We used agarose hydrogel as an *in-situ* model porous material since 0.6% replicates the brain’s diffusivity traits. Therefore, we were able to determine agarose hydrogel properties regarding pore size, permeability, and molecular diffusivity in this material from the current literature. Narayanan et al. and Pernodet et al. have both determined the pore sizes of agarose gels [45], [46]. Pernodet et al. analyzed agarose hydrogels for multiple concentrations by using atomic force microscopy across broad swathes of the gel surfaces. This revealed variations in pore sizes allowing measurements of the average size and distribution of pores

The permeability of agarose hydrogels were measured in multiple studies[44], [47]–[52]. Kosto (2005)[44], Johnson (1996 A)[49], Johnson (1996 B)[48], and Johnston (1999)[50] measured permeability in agarose hydrogels using similar protocols. Kosto (2005) matched the results to the theoretical estimates based on the parallel fiber theory. Consequently, we based our permeability values on results derived by Kosto (2005). Johnson (1996 A) established the protocol for calculating permeability, but did not rigorously verify the resultsUsing these results, we developed a curve comparing agarose concentration to permeability. Upon determining the pore size and permeability estimates of agarose hydrogels, we calculated porosity using the equation developed by Aizawa et al. in Equation 5, the porosity (α), permeability (k_1_), and pore size(d_pore_) are related.

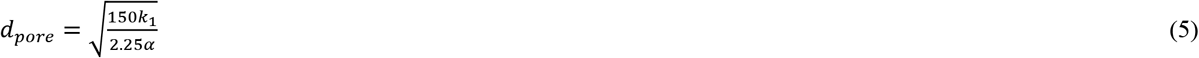

Lead et al gave ranges of diffusion coefficients of different sized molecules in agarose hydrogels relevant to our study[43].

### 3.2 Effects of Output Probe on Diffusion Profile

Previous CED simulations have only utilized a single probe to inject a solution around its outlet. In contrast, our model simulates convection enhanced delivery for targeted flow over a region of interest with the addition of an output probe to add an additional dispersive force on the solute. Consequently, we simulated multiple distances of separation between input to the output probes to determine the effect in comparison to a single injection probe. As hypothesized, addition of an output probe resulted in fluid perturbation when compared to a single input probe (Figure 3). Additionally, we determined that the distance of the output probe from the input probe affected the diffusion profile of the solute

**Figure 3.**
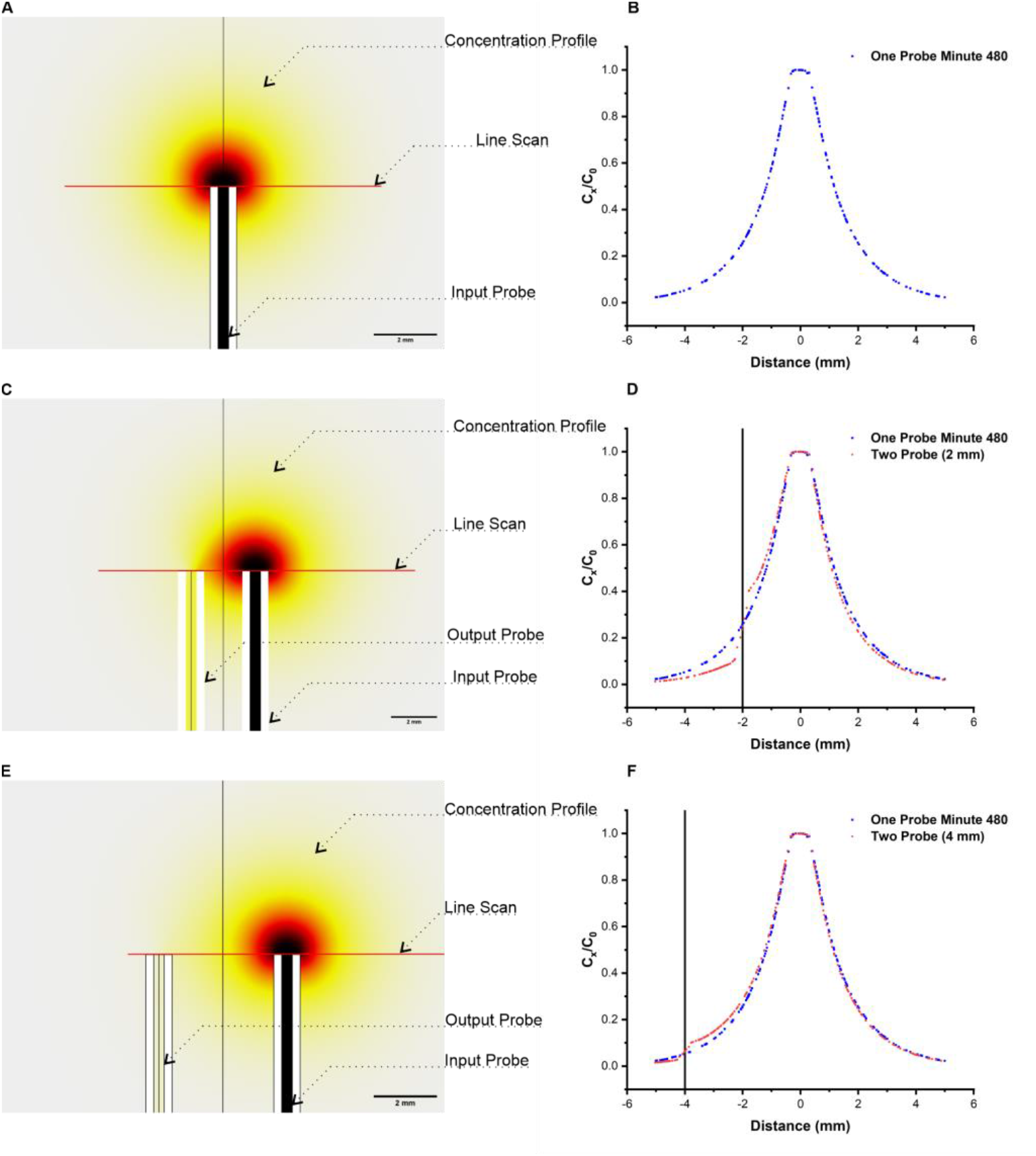
Perturbed Concentration Profiles. Single probe (A and B) illustrates single probe geometry and the single probe diffusion profile resulting from constant injection of a solution over 480 minutes. Two mm probe (BC and D) show a two probe geometry with 2 mm separation resulting in a strongly perturbed profile. Four mm probe (E and F) shows a two probe geometry with 4 mm separation showing decreased but existing impact from convection perturbing the diffusion profile.

Figure 3A and 3B illustrate the diffusion profile of a single input probe via the standard CED technique. Throughout a continuous injection of solute for 480 minutes, the concentration rapidly diminished as the distance from the center of the probe increased, following error function decay. Addition of the output probe 2 mm from the input sampling with an equal output flow induced perturbation in the diffusion profile (Figure 3C and 3D). Between the input and output probe locations, or −2 and 0 mm, on the chart, the concentration was raised relative to the single probe profile—the percent change is shown in Figure S3A. This change was due to increased fluid and solute flow between the two probes that induced a “conveyor belt” effect on particle flow. At distances after −2 mm, there was an induced drop in concentration due to the output probe aspirating the solute free solution from the gel, precluding any further flow of solutes past the probe. Of note, addition of the output probe impacted the diffusion profile on the side contralateral to the output probe placement, inducing a slightly depressed profile when compared to the single probe scheme. In supplemental information, the delta plots of a distance sweep show percent change from the single probe and the modified setup (Fig S3). For the single probe profile and the 2 mm separated probes profile (Fig S3B) the measurements were taken at −1- and 1-mm points on the normalized plot (Figure 3D). After 480 minutes, the Across Input profile distance is depreciated approximately 2.5% while the Near Output profile is increased by nearly 5%. This depreciation on the Across Input side was due to the output probe sampling the solute, thus decreasing its presence elsewhere.

Increasing the separation of the input and output probe by 4 mm from 2 mm diminished the impacts of CED (Fig 3E and 3F). Figure S3 C2, demonstrates that the difference at 2 mm across the input probe from the output probe was nearly 0. The output probe induced a significant delta of approximately 3% that plateaued after 350 minutes, indicating that the system was in steady state. Based on our input conditions, the effective range of impact for CED in our system was a maximum separation of 4 mm between the input and output probes, as evidenced by constant system conditions and a small delta value with increasing probe separation distances. Other systems with differing gel matrices and flowrates will have a different distance effective range. Consequently, with this perturbation, we can direct regions of high and low solute concentration utilizing CED. Translationally, these results suggest that this model could be used to direct high concentration of drugs over the tumor while limiting concentration levels in healthy surrounding tissue in patients undergoing treatment for GBM.

### 3.3 Effect of Parameter Optimization on Diffusion Profile

Our computation model presents a unique opportunity to optimize key variables, most notably the flowrates, flow scheduling, and diffusivity, to selectively deliver solute in a target area via CED As such, we utilized our computational framework in the presence of an output probe to evaluate the impact of adjustments relevant to CED, namely flow rate, solute diffusivity, porous matrix properties, and flow scheduling (Fig. 4). For each parameter, we determined the changes in concentration profiles by calculating the difference (delta) between the profile of the single probe infusion relative to those of the adjusted parameters. This delta value was calculated by subtracting the data points occurring near −3 mm and 3 mm along the left column x-axis plots.

**Figure 4.**
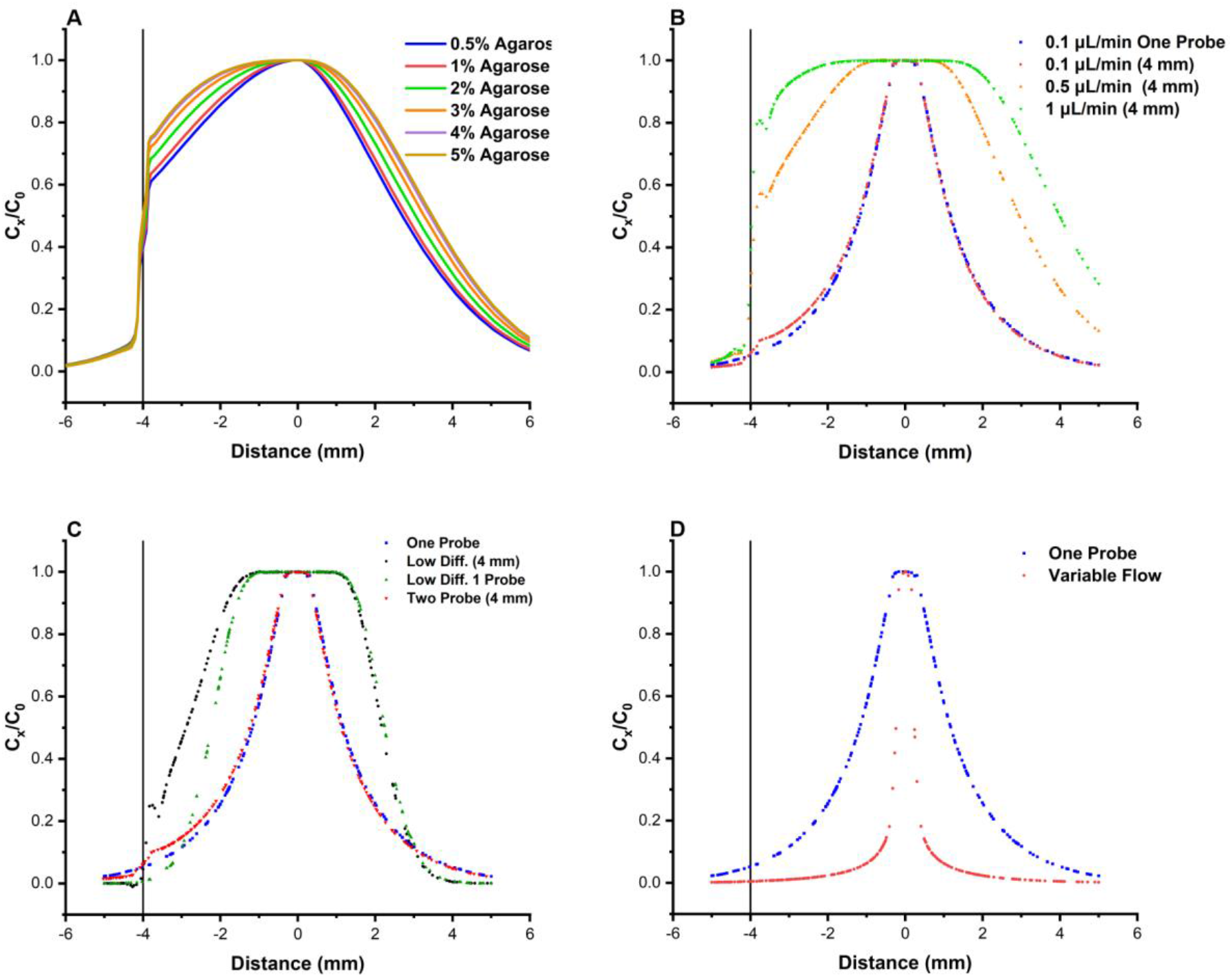
FEM Mechanistic Results. (A) Profile resulting from 2D simulation of porous matrix corresponding to multiple agarose hydrogel concentrations. FEM Optimization of CED. Delta measurements were taken as the difference at −3 mm and 3 mm along the line scans between the single probe profile and the adjusted profile. In the agarose sweep plots (A) there is a decreasing diffusion with increasing matrix density. In the flowrate sweep plot (B) the flowrates are increased from 0.1 μL/min to 0.5 and 1.0 μL/min showing increasing CED impact with increasing flow. The Low Diffusivity plot (C) illustrates the impact of diffusivity of a large biomolecule which results in solute minimally diffusing from the input probe. Variable Flow plots (D) demonstrate variable flow regimes with this simulation utilizing. Over 8 hours the input and output probes oscillated in running every hour.

To model different tissue types, we simulated CED with changing porosity and permeability based upon literature-derived values for each property in agarose hydrogels in a 2D plane [44], [45]. With increasing concentration of agarose polymers, the porous matrix density increases. This increase impeded solute flow, resulting in a greater concentration of the solute near the input probe (Fig. 4A) Modulation of flow rates can also impact solute diffusion by establishing a higher concentration gradient near the input probe as more solute is delivered. We utilized a single probe and 4 mm separated input/output probe scheme perfusing solution at 0.1 μL/min for the input and output (Fig. 4B). The two probe 4 mm scheme was then iterated with an input and output flowrate increase to 0.5 μL/min and 1 μL/min. Increasing the flowrate increased perturbation due to an enhanced rate of particle movement (“conveyor belt”) between the input and output probes.

A solute’s diffusivity, or ability to diffuse in a media, will impact its concentration around the input probe following focal delivery. Higher molecular weight drugs most often have lower diffusivity, impeding their ability to diffuse and permeate through the tumor volume. In Figure 4 C lower diffusivity shows diminishing solute dispersal from diffusion but increased dispersal from momentum transfer induced by the conveyor belt. The high concentration values of the low diffusivity scatter around the probe tip indicating that much of the solute remained around the input probe. However, at low diffusivity, the single versus dual probe design demonstrates that convection can enable motion of low diffusivity molecules based on 35% increase in solute concentration with the addition of an output probe at low diffusivity conditions.

We then wanted to study the effect of variable flow which is used to optimize dosing on tumorous and healthy tissue. For variable flow, the input flow was alternated on and off every other hour for the full 480-minute run; the output probe had an inverse sampling schema. The variable flow paradigm delivered 50% less solute. Our experiments demonstrate that the flow rates, diffusivities, and probe placement have the greatest perturbation relative to the single probe. Also, notable are the time components in that most of the profile adjustments measured by the delta values plateau overtime except for diffusivity.

### 3.4 Comparison with In Situ Trial

In supplemental information Figure S 4, we compared our simulation results from a 4 mm separated probe simulation to in *situ* trials. The experimental trials were conducted four times with the line scan plots averaged to develop error bars. The simulation results of an input and output probe described in Table 1 including for porosity and permeability closely aligned with the in *vitro* results, particularly in regard to the decay profile from −2 mm to the input probe and on the side contralateral to the output probe. The greatest difference between *in vitro* and *in silico* results occurred near the output probe at −4 mm due to optical disruption of the output probe presence.

## 3. Discussion

It is estimated that less than 2% of drugs can permeate the blood-brain barrier, rendering drug delivery into the CNS a significant challenge for therapeutic progress against GBM [55]. CED minimizes drug delivery issues by injection directly into the brain extracellular matrix (ECM) rather than delivery across the BBB[56]. Despite circumventing the BBB, treatments with convection enhanced delivery have not yielded improved results in patient survival[6], [54], [56], [57]. Computational models of drug delivery have the potential to improve CED by empowering optimization of multiple parameters, including catheter placement and flow rates, to maximize the volume of tumor permeated by a candidate therapy.

Few CED studies have sought to optimize parameters such as probe placement, flowrates, and the number of probes to maximize drug permeation within tumor tissue while sparing nearby healthy tissue. Our model is modular in the application of additional probes, tissue variations, and the ability to scale complexity to incorporate or isolate variables. To that end, we simulated a more complex method with multiple probes to induce increased directionality of the concentration profile toward a region of interest. A 6-probe array including two injection (input) probes and four output (extrusion) probes were placed in an agarose hydrogel within a petri dish (Figure 5). Placement of the output probes near the input probes limits solute spread to surrounding healthy tissue, while the other two output probes direct solute flow over the target zone of interest. Solute dispersion, depicted by iso-curves illustrating a curve of constant concentration, was directed over the middle region of interest via CED-induced perturbed diffusion. For GBM treatment, this middle region would include the GBM allowing the conveyor belt of drugs to pass over the GBM while limiting drug off target effects by decreasing drug presence in the surrounding healthy tissue. In therapeutic clinical trials, we foresee this computational modeling streamlining early phase planning. With the knowledge of the pharmaceutical molecular size, the infusion can be computed for a variety of schemes. Once applied, differences between the computational model and trial results can be isolated to remaining factors such as cellular uptake.

**Figure 5.**
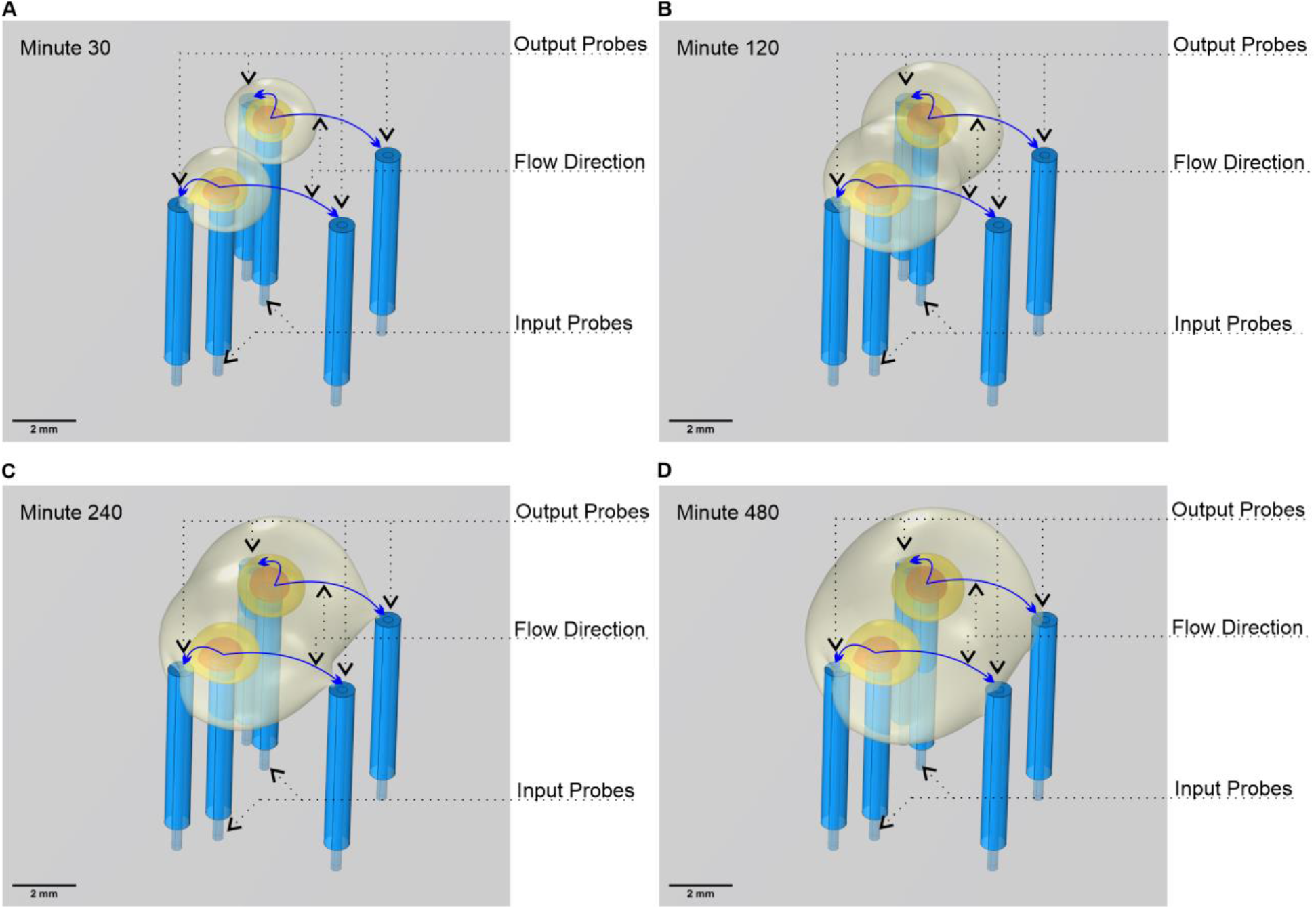
Improved directionality. Six-probe array determined by finite element analysis for improved convective enhanced delivery for an application demonstration. Two probes inject solution into a porous matrix with the other four withdrawing to direct solute dispersal. (A) Solute isoconcentration profile after 30 minutes of delivery. (B) Solute iso-concentration profile after 120 minutes of delivery. (C) Solute iso-concentration profile after 240 minutes of delivery. (D) Solute iso-concentration profile after 480 minutes of delivery.

Precision control of parameters in CED can allow for drug targeting and dispersion within brain tumors. Of all parameters tested in this study, flowrate changes induce substantial impact on the change in the concentration profile relative to baseline. However, fast flowrates, particularly at the input probe, are limited by fluid backflow in the probe if fluid cannot diffuse quickly enough at the injection site. Although a solute’s diffusivity may not be as easily parameterized, we have found that decreasing diffusivity with increasing particle size induced significant changes in the concentration profile for both the single and dual probe methods. Decreased diffusivity resulted in diminished solute diffusion, increasing its concentration in the vicinity of the input probe. These results confirm the importance of a drug’s diffusivity properties when new therapies are developed and when considering new CED methods for improved drug distribution. For Figure 4 C (reduced diffusivity), the plot relays substantial differences between the single probe, decreased diffusivity plot, and single probe with decreased diffusivity. The plot shows limited motion of the low diffusivity solution, but shows substantial change between the outlet and non-outlet side of the inlet probe. This demonstrates the relative increasing contribution of convection in contrast to diffusion. That is, the convection induces large changes. Plots with similar delta values between the output and non-output sides denote limited perturbation from the identified parameter. Whereas, for the variable flow profile, the input solute was less than the other iterations allowing the solute to diffuse more rapidly. While not a trait to optimize, tissue density within infusion regions vary which impacts solute dispersal. We show that for tissue with denser matrices, more of solute stays near the infusion probe due to increase hindrance to diffusion (Fig. 4A). This leaves two factors for optimization with CED. For dense tissue, additional probes may be required to force the solute into a dense region—tumorous advancing regions are usually denser than healthy brain tissue. Whereas, for infusion near a non-dense region, solute may evacuate the zone quickly requiring multiple probes to limit dispersal. An issue with the plotting of the delta values, however, was in direct comparison from the adjusted profile to the control profile. Due to quantized results, the distances from the input probe did not exactly align from plot to plot so we plotted the nearest located values to calculate the delta difference.

## 4. Conclusion

We have demonstrated that convection enhanced delivery in conjunction with an extrusion probe can perturb the diffusion profile of an injection solute, allowing for control of solute diffusion within a porous matrix. Furthermore, optimization of the relevant parameters can further increase the degree of control over solute spread, which may be of interest in limiting the deleterious impacts of therapies on adjacent normal tissue. Furthermore, variations of parameters such as solute diffusivity, flow scheduling, flowrate, and initial concentration can be modulated to change the concentration profile of an injected solute. Optimizing these parameters via computational modeling and then *in situ* within the human tumor could provide an as-of-yet untapped opportunity to improve ongoing CED studies. In conclusion, we conducted FEM simulation of CED in COMSOL Multiphysics to model CED with input and output probes injecting solute within porous matrices simulating hydrogels and varied multiple critical parameters resulting in adjusted concentration profiles. Future work building from this study will aim to simulate the tumor as a material of separate density from the surrounding brain material in its anatomically accurate shape to construct the physical boundary conditions.

## 5. Acknowledgment

MRK and TB conceived and supervised the project. MRK acknowledges the funding support of the VPRI startup fund. We also knowledges the contribution of Lawrence Ray, Braden Norris, and Gerra Licup from the University of Nevada, Reno, for initially collecting experimental data in agarose diffusion experiments. MRK, TB, CS, and CRC contributed to the writing. KZH aided in FEA analyses and analyzing results. MRK and CS designed; CS executed and experimented with the project; analyzed the results; and wrote the Manuscript.

## Supplemental Information

**Figure S1:**
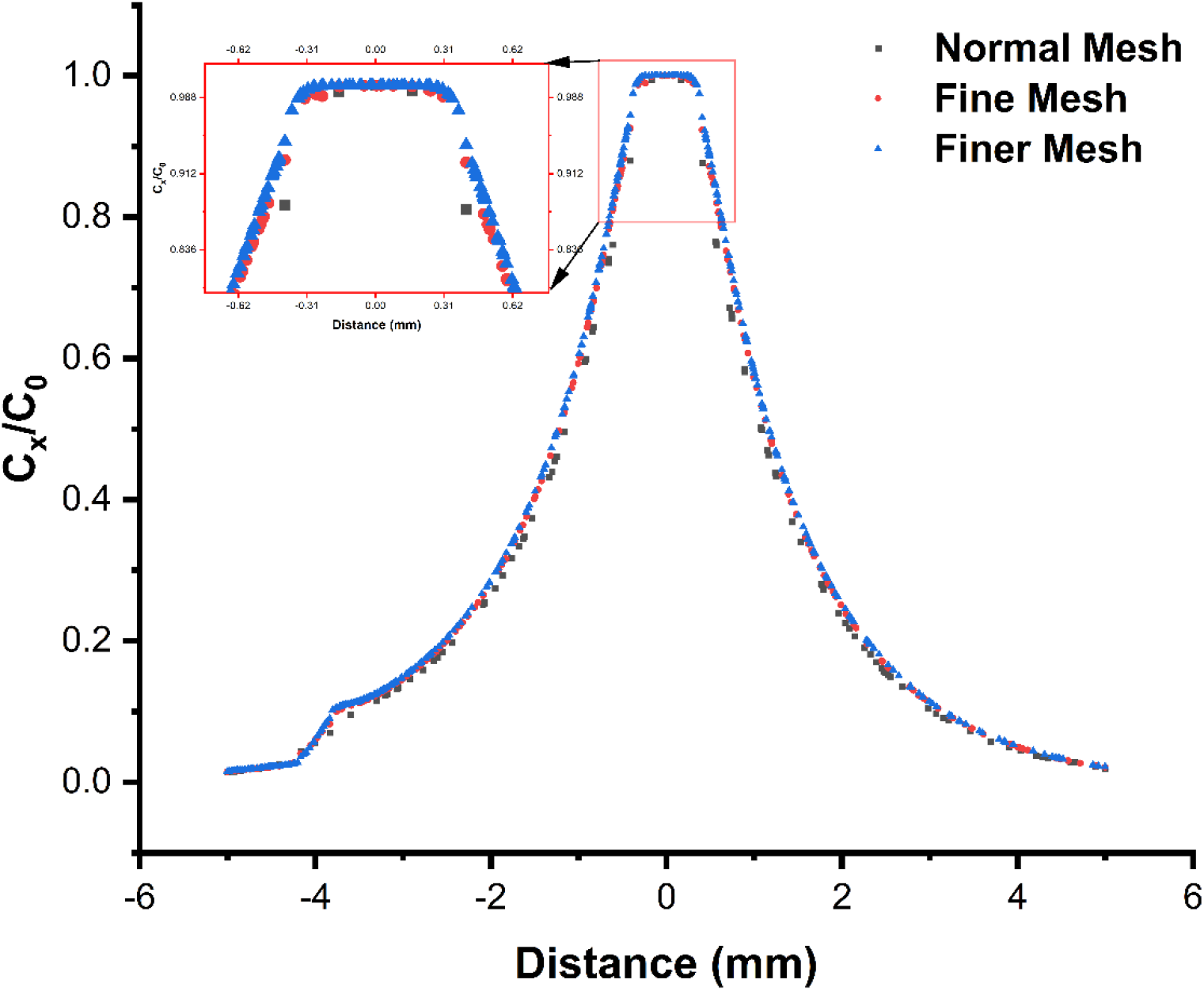
Meshing Convergence. While finite element methods can provide an approximate solution to differential equations, the fineness of the meshing can impact the solution. This figure illustrates the results of three simulations of the 4 mm separated probe setup containing the same geometries and boundary conditions with only the meshing adjusted. Three COMSOL defined meshing sizes being Normal mesh, Fine mesh, and Finer mesh. As can be seen, Normal meshing is slightly different, while Fine and Finer meshing being indistinguishable.

**Table S1.** Meshing characteristics used for the simulations

| Element Type | 2 Probe 4mm apart | 1 Probe | 2 Probe 1mm apart | 2 Probe 2mm apart | 6 Probe |
| --- | --- | --- | --- | --- | --- |
|  | Number of Elements | Number of Elements | Number of Elements | Number of Elements | Number of Elements |
| Tetrahedron | 419314 | 340183 | 355096 | 388530 | 751253 |
| Pyramid | 428 | 214 | 428 | 428 | 1284 |
| Prism | 60572 | 47950 | 58628 | 59360 | 106374 |
| Triangle | 24102 | 26310 | 23412 | 23660 | 40624 |
| Quad | 720 | 612 | 720 | 720 | 1152 |
| Edge Element | 1412 | 1207 | 1412 | 1412 | 3460 |
| Vertex Element | 56 | 44 | 56 | 56 | 152 |
| Maximum Element Size (mm) | 4.21 | 4.21 | 4.21 | 4.21 | 4.21 |
| Minimum Element Size (mm) | 0.104 | 0.104 | 0.104 | 0.104 | 0.104 |
| Study Time | 1 hour 18 minutes | 1 hour 1 minute | 1 hour 11 minutes | 1 hour 20 minutes | 2 hours 45 minutes |

**Figure S2:**
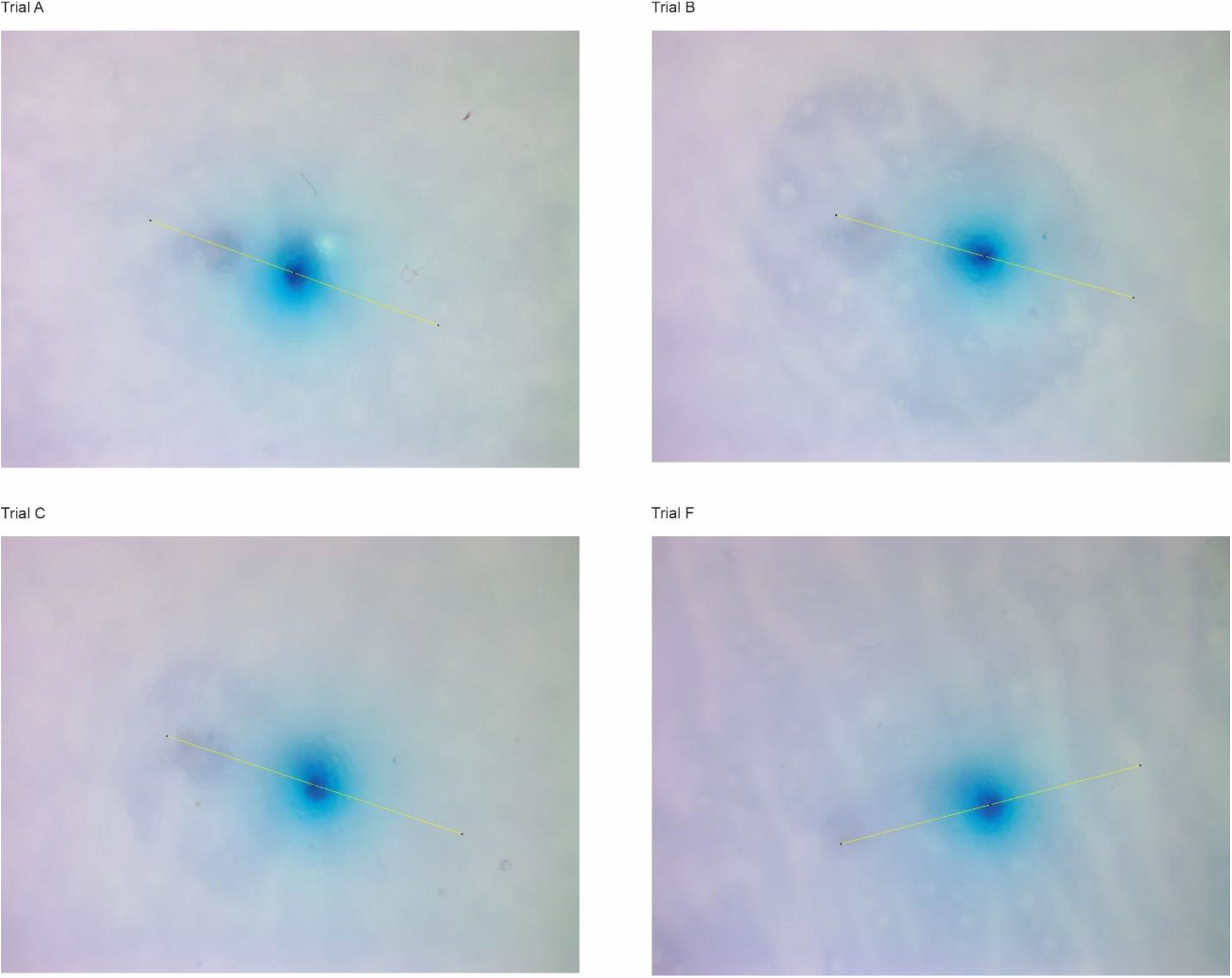
Line scan locations. Experimental convection-enhanced delivery with perturbing output probe image results after 480 minutes.

**Figure S3:**
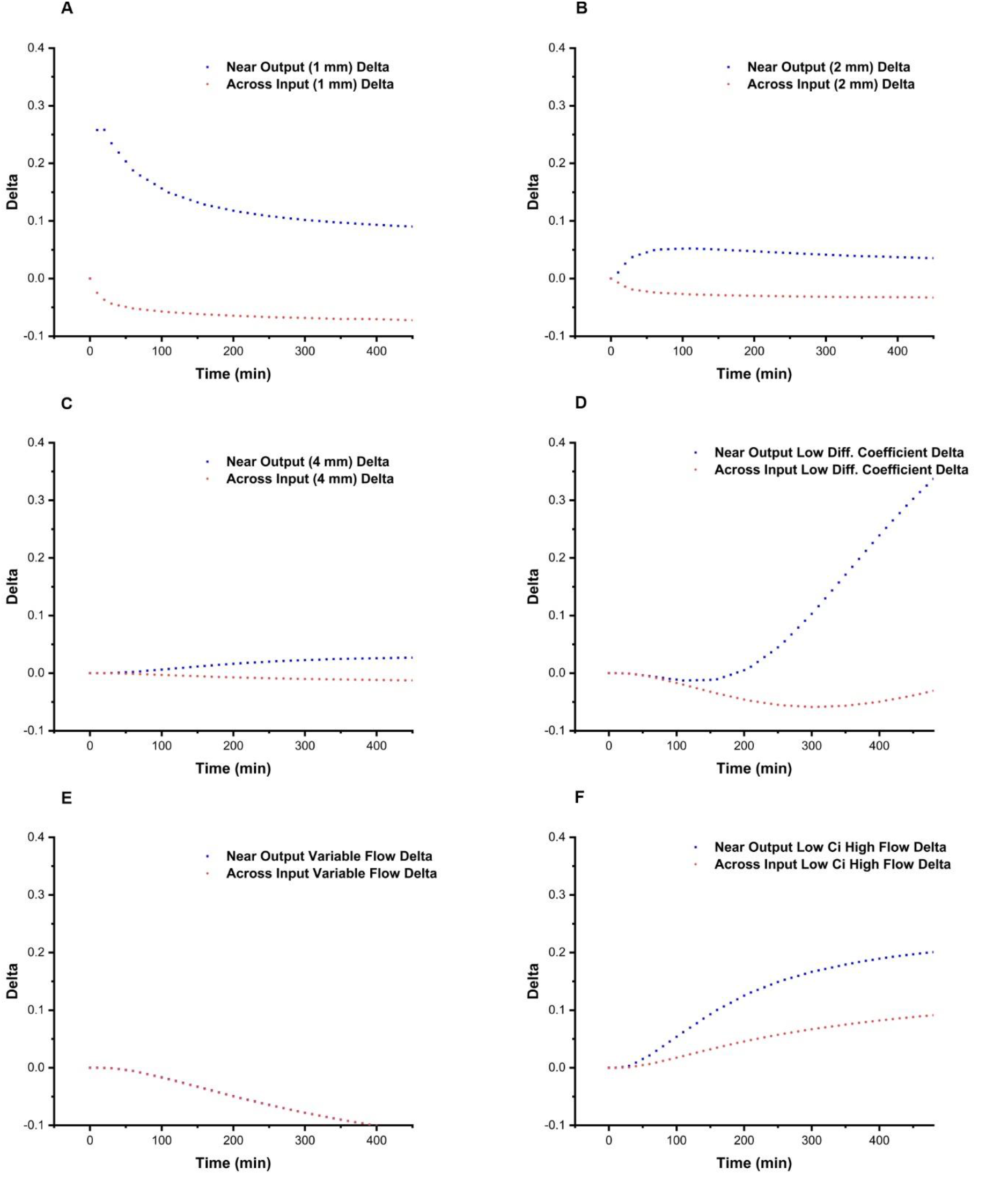
Delta Plots. A. Delta was found at 0.75 mm from input probe. B. Delta was found 1 mm from input probe. C. Delta was found at 3 mm from input probe to complete distance sweep. D. Delta for varying coefficient of diffusion. E. Delta of variable flow simulation. F. Delta for high flow of 0.2 uL/min with reduced concentration to maintain constant mole balance. High concentration values near the input probe indicates that much of the solute remains in close proximity to the input probe. For this situation, this piling up is due to a low concentration gradient not pushing solute away. This indicates greater control of solute dispersal. Due to the piling of solute, the delta values of both sides increase relative to the single probe profile. However, with greater flow, CED greatly increased concentration by nearly 20%.

**Figure S4:**
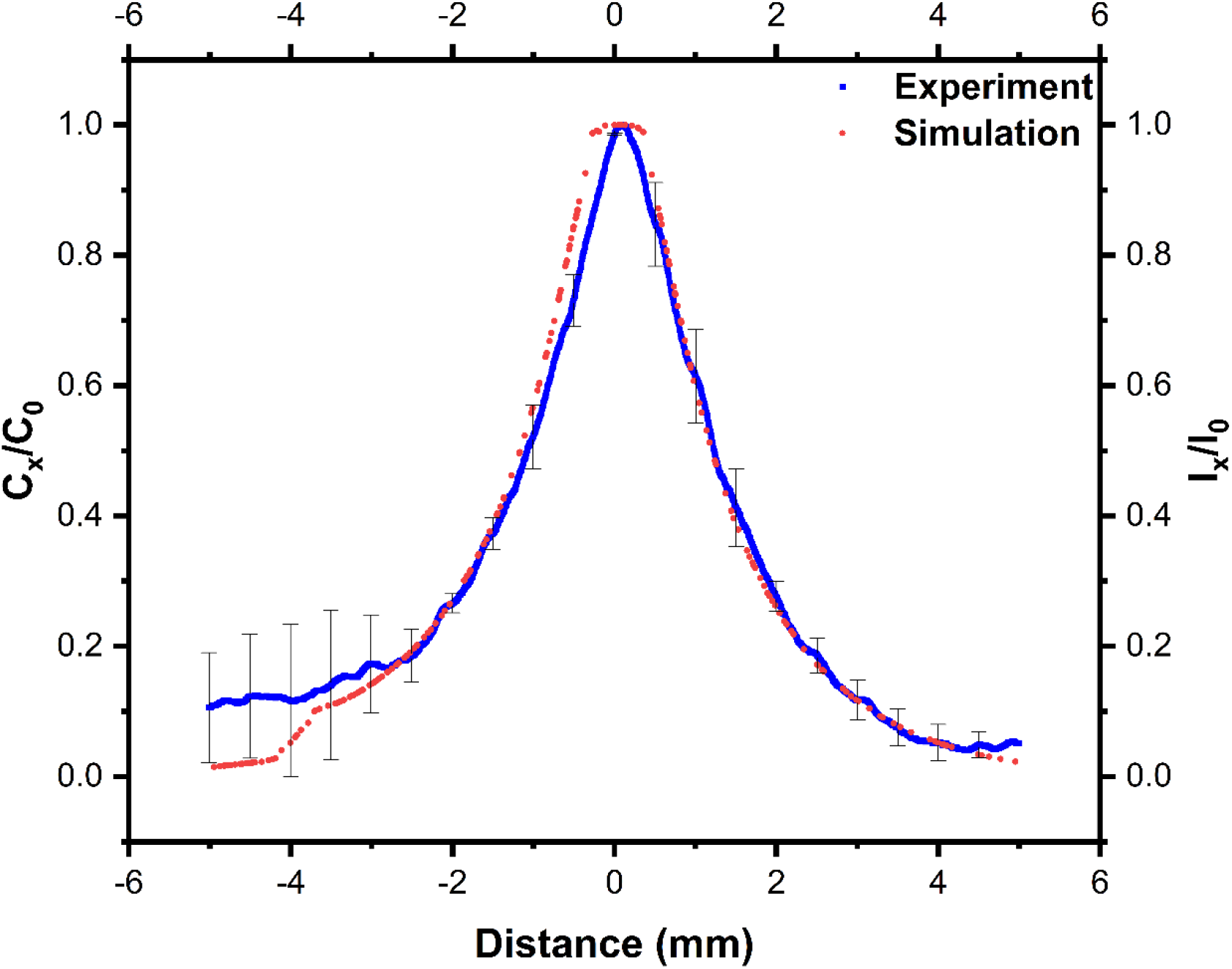
Verification of Porosity and Permeability. Plot comparing the normalized curves of experimental and computational results. Setup includes an input probe located at 0 mm and an output probe at −4 mm. The plot between −4 mm and 0 mm is the region of the conveyor belt and shows increased concentration. Computational model using literature derived values of porosity, permeability, and diffusivity matched the experimental results.

